# Learning the language of protein-protein interactions

**DOI:** 10.1101/2025.03.09.642188

**Authors:** Varun Ullanat, Bowen Jing, Samuel Sledzieski, Bonnie Berger

## Abstract

Protein Language Models (PLMs) trained on large databases of protein sequences have proven effective in modeling protein biology across a wide range of applications. However, while PLMs excel at capturing individual protein properties, they face challenges in natively representing protein–protein interactions (PPIs), which are crucial to understanding cellular processes and disease mechanisms. Here, we introduce MINT, a PLM specifically designed to model sets of interacting proteins in a contextual and scalable manner. Using unsupervised training on a large curated PPI dataset derived from the STRING database, MINT outperforms existing PLMs in diverse tasks relating to protein-protein interactions, including binding affinity prediction and estimation of mutational effects. Beyond these core capabilities, it excels at modeling interactions in complex protein assemblies and surpasses specialized models in antibody-antigen modeling and T cell receptor–epitope binding prediction. MINT’s predictions of mutational impacts on oncogenic PPIs align with experimental studies, and it provides reliable estimates for the potential for cross-neutralization of antibodies against SARS-CoV-2 variants of concern. These findings position MINT as a powerful tool for elucidating complex protein interactions, with significant implications for biomedical research and therapeutic discovery.

## Introduction

The success of large language models (LLMs) in natural language processing—where complex semantic and syntactic relationships are learned from sequences of words—has inspired their application to protein sequences. By treating amino acid sequences as “sentences,” protein language models (PLMs) can implicitly learn structural and functional patterns, enabling predictions of protein folding, mutational effects, and antibody optimization without explicit structural labels [1–9]. However, within cellular environments, proteins rarely act in isolation; instead, they form extensive interaction networks essential for processes such as signal transduction, metabolic pathways, and cellular structural stability. Hence, a comprehensive understanding of protein biology requires moving beyond isolated protein sequences to consider the complex interactions between multiple proteins. Despite this, even PLMs that predict high-resolution structures have thus far been limited to learning patterns from single protein sequences [8, 10].

Since almost every PLM is trained using a self-supervised objective on single chains, existing models often struggle to effectively capture PPIs. Previous approaches that have used PLMs to predict PPIs [11–16] use representations generated from each protein sequence independently without incorporating the contextual information from the interacting partners. This independence causes critical interaction-specific features to be overlooked, as the representations of each protein sequence are generated in isolation. Furthermore, this approach becomes increasingly impractical for PPIs that involve complex multi-sequence (2+) interactions, such as those seen in antibody-antigen or TCR-epitope-MHC complexes [17, 18]. While concatenating input sequences has been proposed as a workaround, it risks degrading embedding quality by treating all sequences as a single unified entity, potentially masking distinct sequence-specific features [19]. To address these challenges, we propose MINT (Multimeric INteraction Transformer), a novel extension of the single-sequence paradigm that allows PLMs to learn distributions of sets of interacting protein sequences. We hypothesize that this approach will enable PLMs to produce context-aware representations that more accurately capture the nuances of protein-protein interactions.

MINT contains two new conceptual advances that enable it to model PPIs effectively. The first is the modification of the popular Masked Language Modeling (MLM) objective so that our model can learn from the interacting protein chains present in STRING-DB, a comprehensive and high-confidence database filtered to contain information on 96 million experimental and predicted PPIs [20]. By training a PLM on this vast corpus of interactions, we aim to uncover deeper representations of not just individual proteins, but also how sets of proteins function together in concert. To the best of our knowledge, no existing PLM has been trained extensively on STRING-DB using a self-supervised pre-training objective. Our second innovation is the adaptation of the model architecture and the training of the popular PLM ESM-2 [10] to handle multiple inputs of protein sequences at once. We present a strategy that allows us to fine-tune ESM-2 on STRING-DB by adding a cross-attention module that explicitly extracts inter-sequence information [21].

We comprehensively benchmarked MINT against widely used PLMs across multiple PPI prediction tasks. MINT consistently outperformed baseline models in binary interaction classification, binding affinity prediction, and mutational impact assessment, achieving a new state-of-the-art AUPRC of 0.69 on the gold-standard dataset constructed by Bernett et al., [22] and delivering a 29% improvement in predicting binding affinity changes upon mutation on the SKEMPI dataset [23]. Further, MINT demonstrated superior performance in antibody modeling, exceeding antibody-specific baselines on property prediction tasks from the Fitness Landscapes for Antibodies (FLAB) benchmark [24] by over 10%. It also outperformed AbMap [25] in predicting binding affinity changes in SARS-CoV-2 antibody mutants, achieving a 14% performance gain in the setting where only 0.5% of the data was available for training. In TCR–epitope modeling, MINT surpassed state-of-the-art models like PISTE [17] and AVIB-TCR [26] with minimal fine-tuning. Finally, we show how MINT can model variant effects in PPIs. In oncogenic PPI (oncoPPI) analysis, MINT effectively distinguished between pathogenic and non-pathogenic interactions, with its predictions matching 23 of 24 previously experimentally validated mutational effects. Finally, in the task of antibody cross-neutralization prediction against SARS-CoV-2 variants, MINT achieved high precision-recall performance, capturing shifts in neutralization profiles across Omicron sub-variants and demonstrating an 80% accuracy in identifying antibodies with consistent neutralization capabilities. By treating multimeric interactions as interdependent sets of sequences rather than isolated pairs, MINT provides a unified approach to computational PPI modeling, offering a powerful framework for studying disease mechanisms, guiding therapeutic design, and advancing immunological research.

## Results

### Overview of the MINT model and training

Protein language models (PLMs) have been applied successfully to predict the structural, functional, and evolutionary attributes of proteins [1, 3–7]. However, encoding protein-protein interactions (PPIs) poses unique challenges, as PLMs are traditionally trained to model single protein sequences independently. To address these challenges, prior approaches have involved passing each sequence through the PLM separately and then concatenating the embeddings or alternatively embedding interacting proteins as a single continuous sequence (Fig. 1a) [16]. However, these methods introduce limitations, such as the loss of inter-residue context and an oversimplified treatment of multi-sequence interactions.

**Figure 1:**
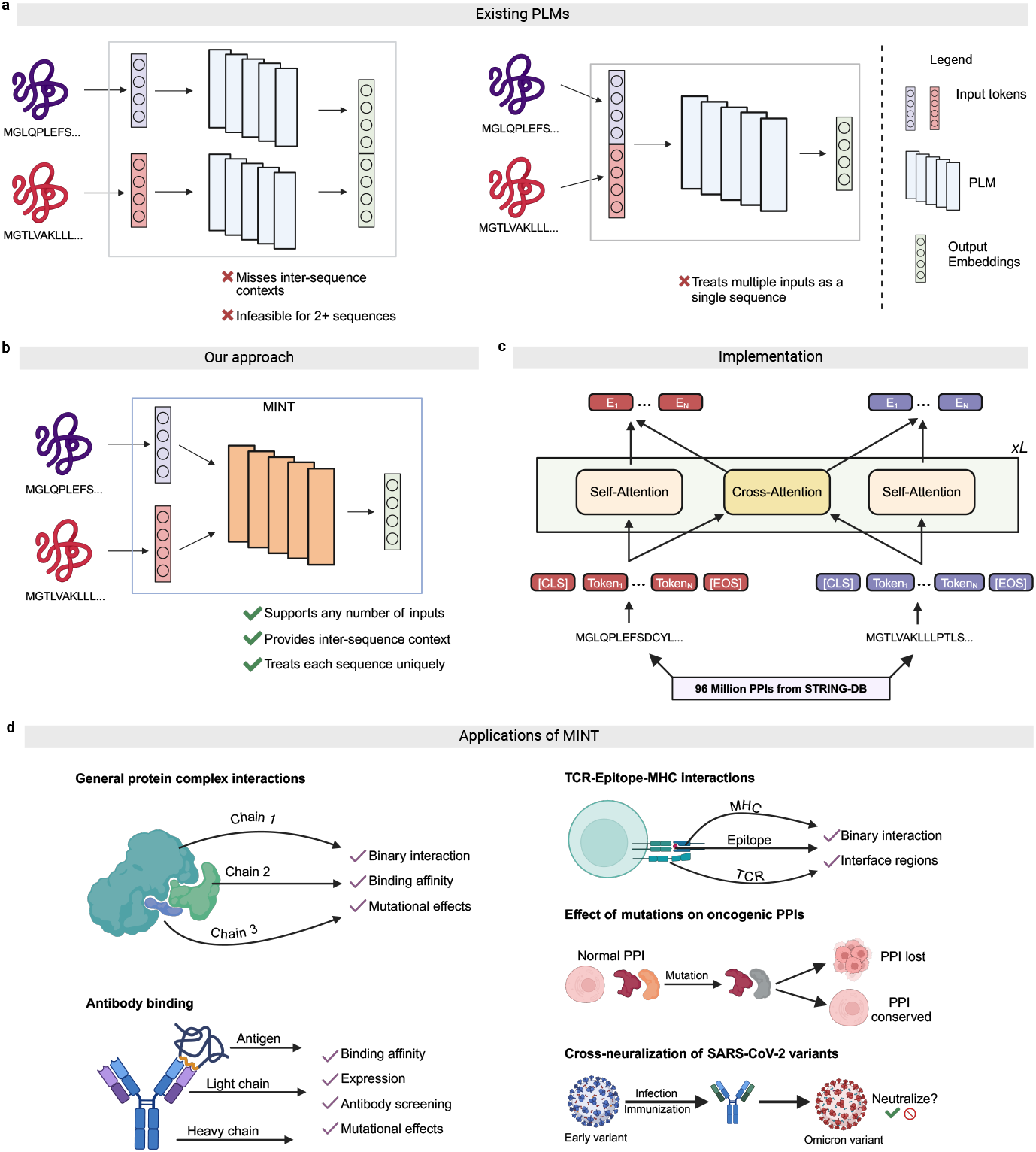
Approaches to PPI modeling and MINT overview. **a**, Existing PLMs either process multiple interacting proteins either by concatenating the output embeddings (left) or concatenating the input tokens (right). The former involves making multiple passes into the PLM to generate embeddings for each sequence independently, and then concatenating them. The latter treats interacting sequences as a single sequence, and generates the embeddings for the concatenated sequence. **b**, MINT treats multiple interacting sequences as separate entities, and generates embeddings in a contextual manner that conserves crosssequence relationships and maintains scalability. This enables it to learn from the vast number for physical PPIs from STRINGDB [20] using the modified version of the MLM loss. **c**, The workflow and architecture of MINT. Each protein sequence is tokenized using the ESM-2 tokenizer [10] and special tokens are added for the start and end of sequence. Note here that we add these special tokens for each interacting sequence, maintaining sequence identity. Our architecture involves the adding of cross-attention blocks to the base ESM-2 model. This results in the output representations of each token being affected by the tokens in the same sequence, and those in the interacting sequences. Each block is repeated *L* times, where *L* is 33 for MINT. **d**, A non-exhaustive list of protein types, PPI properties and research questions that can be evaluated using MINT. We benchmark it on general protein complex, antibody and TCR-Epitope-MHC interactions against other PLMs. We then give examples of the type of novel analysis that can be done using MINT by predicting oncogenic PPIs and SARS-CoV-2 antibody cross-neutralization using experimentally labeled data.

To address these limitations, we developed MINT, a PLM specifically designed to model PPIs by enabling the simultaneous input of multiple interacting protein sequences (Fig. 1b). Building on the 650-million parameter ESM-2 architecture, MINT introduces a novel crosschain attention mechanism that preserves inter-sequence relationships and scales effectively to handle complex interactions with more than two chains. Whereas the self-attention mechanism in ESM-2 utilizes rotary positional encoding to capture intra-sequence positional relationships, MINT applies self-attention solely within chains and incorporates additional attention blocks for cross-chain interactions without rotary encoding [10]. This modification ensures that each token representation in our model captures contextual information across all input sequences (Fig. 1c). Further architectural details and pseudocode are provided in Methods.

To train MINT, we curated a dataset from the STRING database, which includes 2.4 billion physical PPIs and 59.3 million unique protein sequences. By applying clustering and diversity measures, we refined this dataset to 96 million high-quality PPIs involving 16.4 million unique sequences (Methods Sec 1). This dataset serves as the foundation for training MINT using a masked language modeling (MLM) objective augmented with cross-attention [27]. Unlike traditional MLM, where token prediction is conditioned solely on intra-sequence context, MINT leverages cross-chain representations to capture co-evolutionary constraints imposed by interacting residues (Fig. 1c, Methods 2). Model training followed a multiphase approach, beginning with the initialization of attention and embedding weights from ESM-2. This warmstart approach allowed us to preserve foundational sequence-level knowledge while optimizing for cross-chain interactions.

The model was trained with a masking scheme and hyperparameters closely aligned with ESM-2, with the objective function incorporating both sequence-specific and interaction-specific signals (Methods Sec 3).

Performance during training was evaluated using perplexity on a validation set derived from a random split of the curated dataset (Supplementary Fig. S3). Together, these architectural and training innovations enable MINT to overcome the inherent limitations of existing PLMs in learning PPI rules. We demonstrate versatility in handling diverse protein sequence inputs, including general complexes, antibody heavy and light chains, T-Cell Receptor (TCR) regions, and peptides, without restrictions on the number of sequences processed concurrently (Fig. 1d). This capability allows MINT to capture intricate features relevant to disease-specific interactions and mechanisms, such as mutational impacts in cancer and the cross-neutralization potential of SARS-CoV-2 antibodies.

### Comprehensive benchmarking of MINT on PPI prediction

PPI prediction is fundamental to understanding cellular processes, offering insights into disease mechanisms and therapeutic targets. Although structure-based models such as AlphaFold [7] provide detailed predictions, they face challenges with scalability and accuracy for non-interacting pairs [15]. Sequence-based methods, in contrast, offer efficiency and flexibility but fall short in explicitly encoding PPIs. To address these gaps, we benchmarked MINT against widely used PLMs in various supervised tasks, including binary interaction classification, binding affinity prediction, and mutational impact assessment.

We initially assessed the performance of MINT in predicting the presence or absence of interactions between protein chains, or binary interaction prediction. This evaluation was conducted using three distinct datasets, including a comprehensive dataset curated by Bernett et al. [22] that employs rigorous sequence similarity thresholds to mitigate data leakage, alongside two additional datasets derived from the PEER benchmark, which specifically focus on binary predictions of PPIs present in human and yeast systems [12]. Next, we evaluated how well MINT can predict absolute binding affinity by leveraging binding data for protein–protein complexes from PDB-Bind, where each complex can have a variable number of protein chains. [28]. Both the binary interaction prediction and binding affinity prediction tasks involve a single interacting group of protein sequences, and an overview of the prediction process is illustrated in Fig. 2a.

**Figure 2:**
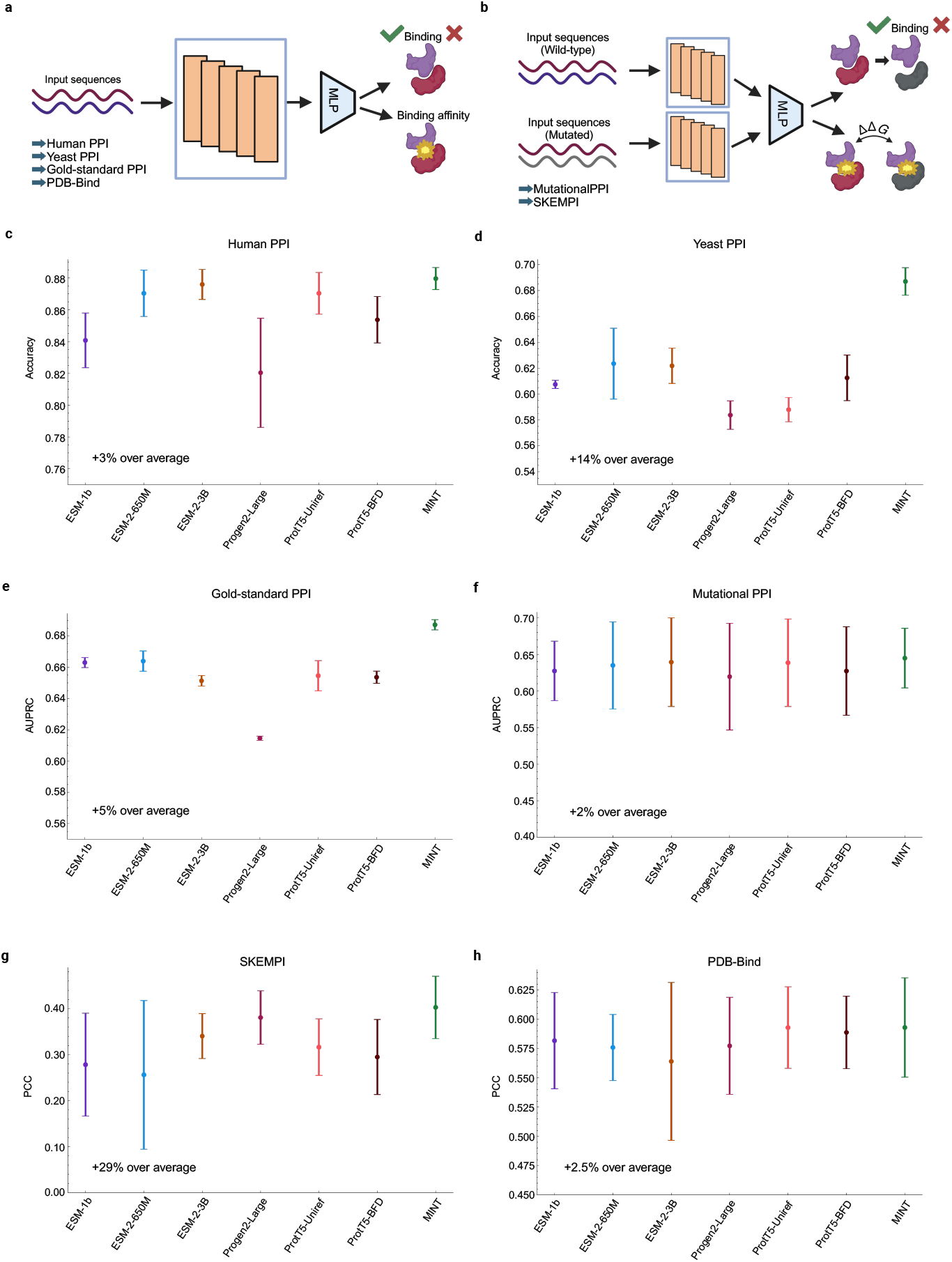
Performance of MINT versus other PLMs on general PPI tasks. **a**, Overview of the model framework for downstream tasks involving the prediction of properties for a single interacting group of sequences. This involves the goldstandard dataset (Gold-standard PPI) [22], the human (HumanPPI) and yeast (YeastPPI) datasets from the PEER benchmark [12, 33] and the PDB-Bind affinity (PDB-Bind) dataset [28]. We generate embeddings for the baseline PLMs and MINT and use a Multi-Layer Perceptron (MLP) to predict the either the binary interaction or the binding affinity. **b**, Overview of the model framework for downstream tasks involving the prediction of properties for mutation effect analysis, involving wild-type and mutated sequence groups. This involves the datasets from SKEMPI [23] and the binary prediction of PPI binding after mutation (MutationalPPI) [29]. We generate embeddings for the baseline PLMs and MINT for both wild-type and mutant sequence groups separately, and then aggregate them to get a final embedding. Then, we use an MLP to predict the either the binary interaction or the change in binding affinity. **c-h**, Results for all 6benchmarking tasks against baseline models with 3 experimental repeats each **c**, Human PPI, **d**, Yeast PPI, **e**, Gold-standard PPI, **f**, Mutational PPI, **g**, SKEMPI, **h**, PDB-Bind. AUPRC: area under precision-recall curve; PCC: pearson correlation coefficient.

We also evaluated the ability of MINT to resolve mutational effects on binding using two datasets. The first contained data from SKEMPI—a database documenting changes in binding affinity due to mutations in one of the interacting proteins—we evaluated the models with previously established cross-validation splits by protein complex [23]. The second task in this context involved predicting whether two human proteins remain bound after a mutation occurs in one of them, thereby testing the model’s sensitivity to sequence alterations that impact binding outcomes [29]. These tasks involve encoding the wild-type protein pairs and the mutant protein pairs, as described in Fig 2a.

In each task, MINT ‘s embeddings were utilized to train lightweight predictor models, facilitating equitable comparisons with baseline pre-trained language models (PLMs) such as ESM-1b [30], ESM-2 (comprising both 650M and 3B parameters) [10], ProGen (3B parameters) [31], and ProtT5 (3B parameters, specifically trained on the UniRef and BFD databases) [32] (Methods section 4.1). We compared two baseline embedding strategies: (1) concatenating independently computed embeddings from the PLM and (2) concatenating interacting sequences as a single input to the PLM (Fig. 1). For all tasks except PDB-Bind, the first strategy of concatenating embeddings worked better for all baseline models.

For the PDB-Bind task, we evaluated only the second method of concatenation, since each input can contain a variable number of interacting sequences, making it impractical to perform a variable number of PLM calls. We show the results for the best performing embedding strategy in Fig. 2c. Results for both embedding strategies are available in the Supplementary Table 1.

MINT consistently outperformed all baseline PLMs across all tasks (Fig. 2c). On the gold-standard dataset constructed by Bernett et al., MINT achieved a new state-of-the-art performance with an area under the precision–recall curve (AUPRC) of 0.69, demonstrating its ability to excel under challenging conditions and to capture non-trivial interaction signals missed by traditional PLMs. In the other binary PPI prediction tasks, MINT achieved an improvement of approximately 14% over the average baseline for yeast interactions and 3% for human interactions. On the SKEMPI dataset for predicting binding affinity changes upon mutation, our model achieved a 29% improvement, representing the highest performance for any model using only sequence information. Notably, MINT consistently outperformed the base ESM-2 model, indicating that the architectural enhancements and fine-tuning on STRING-DB enable the learning of essential PPI signatures absent in ESM-2. Furthermore, its performance advantages over larger, general-purpose PLMs like ProGen and ProtT5 underscore the importance of task-specific adaptations over mere increases in model size.

### Performance on antibody modeling tasks

Building on its superior performance in general PPI tasks, we extended MINT to domain-specific challenges in antibody and immune modeling to test its ability to generalize beyond traditional protein interaction datasets. Antibodies are essential for the adaptive immune system’s ability to recognize and neutralize pathogens. They are symmetrical, Y-shaped molecules composed of two heavy chains and two light chains [34]. Antibody-antigen interactions are also uniquely difficult to model because CDR regions are highly variable and not as constrained by evolution [35]. Here, we show how MINT can be used to predict the binding characteristics of antibodies to antigens by leveraging its ability to jointly model the heavy and light chains, enabling richer representations suitable for various downstream tasks like binary interaction and binding affinity prediction. We compare MINT with three deep-learning-based approaches specific to antibody modeling, evaluating our model on the same datasets and employing the same evaluation strategies used in each approach.

We compared MINT against IgBert and IgT5—two models fine-tuned on extensive datasets comprising two billion unpaired and two million paired sequences of antibody light and heavy chains [18]. These models explicitly learn the language of antibodies, encoding heavy and light chains either jointly as pairs or separately in an unpaired manner. We assessed their performance on four supervised property prediction tasks using the FLAB dataset, which includes experimentally measured binding energy data from Shanehsazzadeh et al. [36], Warszawski et al. [37], Koenig et al. [38], and expression data from the latter study. To ensure a faithful comparison, we followed the 10-fold cross-validation procedure outlined by Kenlay et al. [18], fitting a ridge regression model on embeddings from MINT (Fig. 3a, (Methods Sec 4.2)). Performance was evaluated by measuring the coefficient of determination (*R*^2^) between predicted and actual values. MINT achieves a performance boost over the antibody-specific baselines on all 4 datasets, with an increase of over 10% on three of them (Fig. 3b). Moreover, it is able to perform equally well on predictions of antibody binding affinity and expression tasks. This performance gain demonstrates that pretraining on a diverse corpus of PPI data enhances generalizability to antibody-specific tasks.

**Figure 3:**
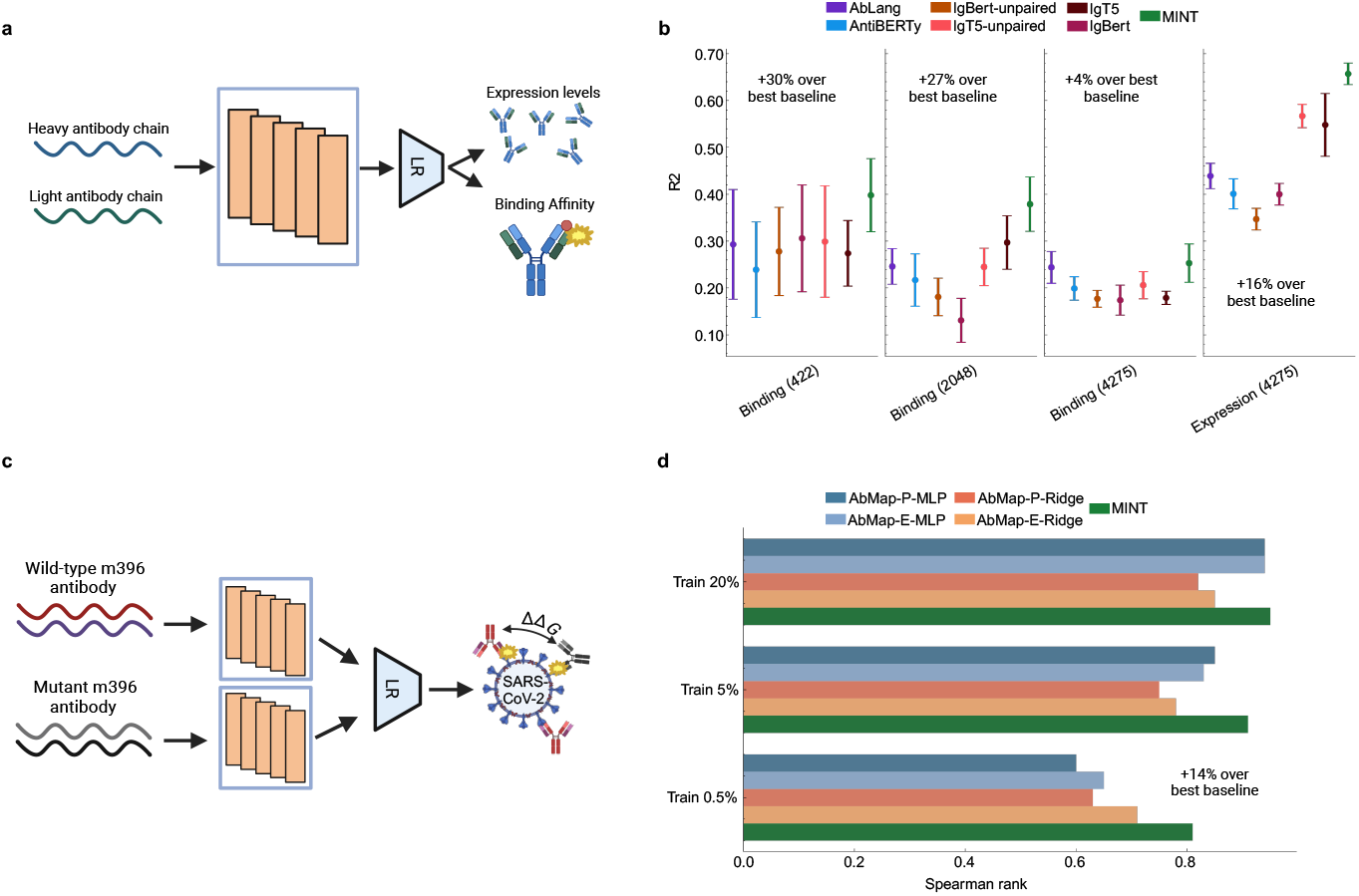
Comparing MINT to antibody-specifc PLMs. **a**, Overview of the model framework for downstream tasks from the FLAB benchmark for evaluating model performance on antibody sequences [24]. MINT treats the heavy and light chains as separate sequences, enabling better representations. We follow previous work in embedding the antibody sequences and training a Linear Regression (LR) model under a 10-fold cross-validation setting to predict either the expression levels or the binding affinity values [18]. **b**, Results for the antibody-specific PLMs and MINT across all four datasets evaluated in the FLAB work. R2: Coefficient of determination. **c**, Overview of the model framework the the prediction of binding affinity changes for m396 antibody mutants against the SARS-CoV-2 virus [39]. We follow previous work (AbMap) in embedding the antibody sequences and training a Linear Regression (LR) model under different fractions of the dataset and testing on the rest [25]. **d**, Results for different configurations of the AbMap model and MINT across all dataset splits evaluated in this work. AbMap-E refers to AbMap with the ESM-1b model and AbMap-P refers to the ProtBert version. The two AbMap models are also evaluated using MLP and ridge regression models for prediction [19].

We further evaluated the performance of MINT in comparison with AbMap, a transfer learning framework fine-tuned specifically for antibodies by focusing on their hypervariable regions and trained on antibody-specific structural and binding specificity datasets [25]. Both models were assessed on a mutational variation prediction task involving the prediction of changes in binding affinity (ddG) for m396 antibody mutants against the SARS-CoV-2 virus, using experimental data from Desautels et al. [39] (Fig. 3c). The m396 antibody originally targets the receptor-binding domain (RBD) of the SARS-CoV-1 spike protein [40]; mutants were generated to evaluate potential binding to the SARS-CoV-2 RBD. Approximately 90,000 mutants were generated in silico, with binding efficacy estimated through ddG scores computed using five energy functions: FoldX Whole, FoldX Interface Only [41], Statium [42], Rosetta Flex, and Rosetta Total Energy [43]. Adhering to the dataset splitting strategy employed by the AbMap method, we trained on designated subsets of the data and evaluated performance on the remaining portions using the Spearman rank correlation (Methods, Sec 4.2). Again, MINT achieves comparable or better performance compared to AbMap over all data splitting instances (Fig. 3d). Crucially, our model achieves a 14% increase in performance when trained on just 0.5% of the samples, suggesting that it can be used for in-silico antibody design even with a small number of training data.

### Learning the language of TCR-Epitope-MHC interactions with minimal finetuning

The interactions between T cell receptors (TCRs), epitopes, and major histocompatibility complex (MHC) molecules are central to the human immune response, allowing the immune system to identify and respond to pathogens, cancer cells, and other threats. TCRs on the surface of T cells recognize specific antigenic peptides (epitopes) that are presented by MHC molecules on antigen-presenting cells (APCs). This recognition is the first step in activating T cells, which is critical for orchestrating an immune response [44]. Several deep learning-based approaches have been developed to model the interactions between TCRs, epitopes, and MHC molecules, primarily focusing on binary predictions of TCR–epitope or MHC–epitope binding (first-order interactions) [45]. More recently, methods have been designed to consider all three entities simultaneously, enabling the prediction of TCR–epitope–MHC binding (secondorder interactions) [17]. Additionally, some techniques aim to enhance specificity by predicting the interface residues between interacting TCR–epitope pairs [46]. MINT offers a flexible framework to model these tasks by accommodating the various input sequences. However, the short length of epitopes and the common practice of considering only the Complementarity-determining region 3 (CDR3) sequences of MHC and TCRs in most datasets present challenges for its direct application, since MINT is not trained on peptides. To address these limitations, we minimally fine-tuned the final layer of MINT to learn TCR–epitope–MHC interactions across the three task types mentioned above (Supplementary Note 3).

For the first-order tasks, we use the TCR-epitope binding prediction benchmark from Therapeutics Data Commons (TDC), which focuses on predicting whether a TCR binds to an epitope [45]. The interaction data is originally sourced from the IEDB [47], VDJdb [48], and McPAS-TCR [49] databases. We compare to popular TCR-Epitope prediction models, namely AVIB-TCR [26], MIX-TPI [50], Net-TCR2 [51], PanPep [52], TEINet [53] and TITAN [54]. We only use TCR CDR3-beta and epitope sequences as input to MINT to maintain a fair comparison with all baseline models (Fig. 4a). MINT obtains an average AUROC score of 0.581, beating the best baseline with an AUROC of 0.576 (Fig. 4b). These results demonstrate that minimal fine-tuning of MINT can effectively model TCR–epitope interactions, even for protein types not encountered during its pre-training, offering a novel approach to improving predictive performance in this domain.

**Figure 4:**
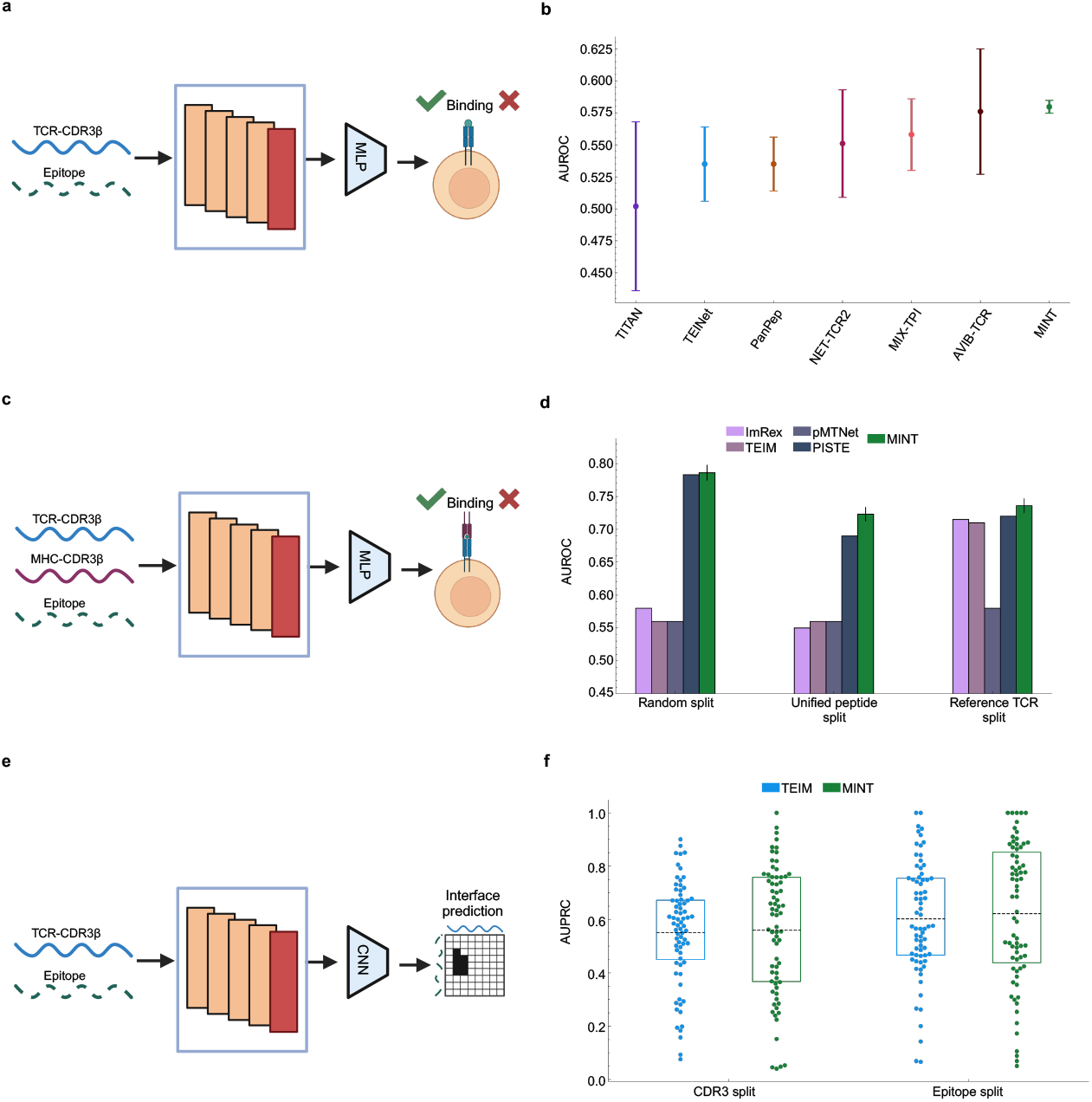
Comparing finetuned MINT to TCR-MHC-Epitope models. **a**, Overview of the first-order interaction prediction task involving TCR-CDR3 and epitope sequences as input [45]. Since MINT has not been trained on short sequences, we finetune its last layer (red block in the architecture) to capture the language of TCR-epitope interactions. We then apply a MLP model to predict whether binding happens or not. **b**, Performance of MINT for predicting binary binding compared to popular TCR-epitope specific models. Results for all baseline models are taken from [45]. AUROC = area under the receiver operating characteristic curve **c**, Overview of the second-order interaction prediction task involving TCR-CDR3, HLA-CDR3 and epitope sequences as input. Again, we finetune the last layer of MINT (red block in the architecture) to capture the language of TCR-epitope-HLA interactions. We then apply a MLP model to predict whether binding happens or not. **d**, Results on the second-order binary binding predictions across different dataset splits on the cancer dataset used for evaluation. Results for all baseline models are taken from [17]. **e**, Visualization of the TCR-epitope interface prediction task using MINT. We feed in the TCR-CDR3 and epitope sequences, embed them using MINT and train a downstream Convolutional Neural Network (CNN) model to predict the interface using a contact map. **f**, Results for the TCR-epitope interface prediction task. We calculate the AUPRC values between the predicted and the true flattened contact maps for each data point in the test set and plot the distribution for MINT and TEIM [46]. This is done across the two dataset splits that we analyzed-test set with unseen CDR3s and test set with unseen epitopes. AUPRC = area under the precision-recall curve

For the second-order interaction prediction, we train our model on the dataset of human TCR, Epitope and Human leukocyte antigen (HLA) interactions, where HLA is the human version of MHCs (Fig. 4c). This binding data was originally procured from the McPAS-TCR, VDJdb and pMTnet [55] databases and curated in the the PISTE model work [17]. The evaluation set used consists of TCR-Epitope-HLA triplets obtained from studies of different cancer types. We used all three dataset splitting schemes as described in the same work and show the performance of the finetuned MINT with PISTE, pMTnet and other first-order models. MINT achieves the highest performance across all dataset splits by leveraging its ability to process multiple protein chains (Fig. 4d). These results highlight MINT ‘s ability to process multiple protein chains and achieve state-of-the-art performance on complex multi-protein interaction tasks. The scalability of MINT to handle an increasing number of protein chains further emphasizes its utility for modeling intricate biological systems.

Finally, for the interface prediction task, we used the dataset constructed in the TEIM work [46], with the TCR-Epitope structure data originally sourced from STCRDab [56] (Fig. 4e).

We restrict ourselves to the splitting strategies where the test set has novel epitopes or novel TCRs that are not seen during training, and each split results in a total of 122 total data points. MINT matches the performance of TEIM on the unseen TCR-CDR3 split, and is better on the unseen epitope split (Fig. 4f). These results underscore MINT ‘s capacity to infer structural relationships between TCRs and epitopes without extensive pretraining and its robust performance with small datasets, suggesting that its internal protein sequence representations are readily transferable to specialized tasks.

### Leveraging MINT for predicting mutation-induced perturbations in oncogenic PPIs

After establishing the superior performance of MINT against popular PLMs on general PPI prediction tasks and against domain-specific models on antibody and TCR-Epitope-MHC tasks, we highlight types of novel analysis that can be achieved with MINT. We first applied it to estimate the pathogenicity of missense mutations affecting PPIs in cancer, focusing on cancer-associated mutations that disrupt PPIs, which we refer to as oncoPPIs. Disease-associated germline and somatic mutations are enriched at PPI interfaces, indicating their potential role in disrupting PPI networks. Cheng et al. [57] identified 470 potential oncoPPIs linked to patient survival and drug responses, and experimentally validated the perturbation effects of 24 somatic missense mutations across 13 oncoPPIs using Yeast Two-Hybrid (Y2H) assays, hypothesizing their functional impact on cancer progression. Existing computational approaches distinguishing pathogenic from non-pathogenic interactions typically focus on single protein sequences, employing biophysical and evolutionary constraints or leveraging protein embeddings (e.g., EVE [58], AlphaMissense [59], ESM-1b [3]). However, in oncoPPIs or any other pathogenic PPIs, mutations rarely significantly affect protein structure but alter protein–partner binding [60, 61]. Therefore, we took advantage of MINT’s ability to effectively model multiple interacting proteins and trained it on a dataset containing information on whether a mutation in a human protein sequence disrupts its binding to another human protein or not [29]. We then predicted the mutational effects on the experimentally validated oncoPPIs from Cheng et al. as outlined in Fig. 5a. Since the training dataset is quite small, we performed multiple repeats of the training runs, or model ensembling, and computed a binding score that is representative of the proportion of times that the model predicts binding over all repeats (Methods Sec 5).

**Figure 5:**
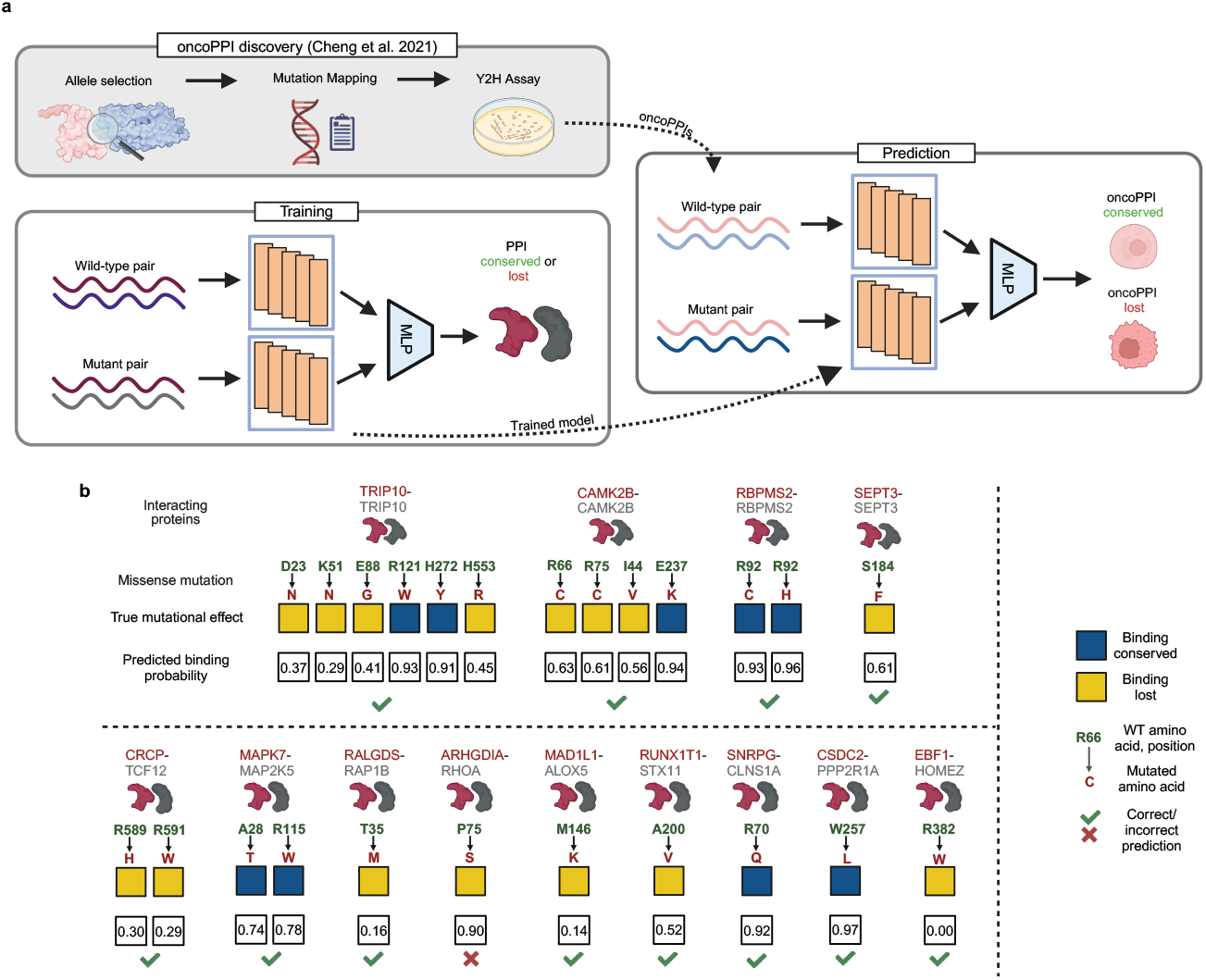
Overview and results of the mutational effect prediction in oncoPPIs. **a**, Outline of the analysis to predict the experimentally-validated mutational effects on binding in oncoPPIs. The oncoPPI discovery was conducted by Cheng et al. [57] to identify 24 mutational effects in 13 different oncoPPIs through computational predictions and experimental Y2H assays. We used MINT embeddings and a trainable MLP on a dataset containing wild-type and mutant human protein pairs to predict whether the mutation results in a loss of binding or not in a fashion similar to Fig 2b. We then used the trained model to predict on the oncoPPI dataset across the 24 mutational effects. Since the training dataset is small, we use an ensemble of 100 trained models and using their consensus to compute the predicted binding score, which is the fraction of times a particular oncoPPI was assigned to be 1 (binding conserved) or 0 (binding lost). **b**, Predicted binding score and true mutational effects from Y2H experiments for all 24 mutations. We show the gene names of the interacting proteins, the missense mutation information (wildtype amino acid and position of the mutation n green and the mutated amino acid in red). Some oncoPPIs are homodimers (similar shaped dark red and gray protein icons) and some are heterodimers (different shaped dark red and gray protein icons). We use green check marks and red crosses to indicate whether our predicted binding score matches the true mutational effect. Using a threshold of 0.68 (Methods Sec. 5), MINT correctly distinguishes 23 out of 24 mutations.

We show the real mutational effects and model predicted scores for all of the experimentally validated oncoPPIs from Cheng et al. in Fig. 5c. Using a threshold value computed in an unsupervised manner (Methods Sec. 5), MINT’s predictions match the mutational effects for 23 out of 24 PPIs. Our model is able to correctly assign the ALOX5-MAD1L1 interaction to be lost after the M146K mutation on ALOX5, and both these proteins have been implicated in tumor progression in several types of cancer [62]. Interestingly, our analysis correctly predicts a destabilizing mutation for the HOMEZ–EBF1 interaction, and previous ZDOCK analysis has shown that the mutation occurs at the binding interface, potentially disrupting the salt bridge and hydrogen bond at that position [13]. Out of all 24 mutations, only one oncoPPI, the ARHGDIA-RHOA pair, was incorrectly assigned. Our results indicate that MINT has the ability to effectively model mutational effects in oncoPPIs and can be used to provide insights into how mutations disrupt PPIs in cancer.

### MINT predicts antibody cross-neutralization against SARS-CoV-2 variants

Next, we consider the task of estimating the cross-neutralization of antibodies against different SARS-CoV-2 variants. SARS-CoV-2 undergoes a very high rate of mutation, leading to a significant change in transmissibility, immune evasion, and severity between strains. One highly mutated version of the virus is the Omicron variant, which was able to evade immunity from several neutralizing antibodies produced by natural infection and vaccination from previous variants [63]. Using experimental methods like deep mutational scans (DMS) to predict immune escape for each newly emerging variant that appears, like Omicron, is time-consuming, expensive, and intractable. Computational methods such as EVEScape, which use pre-pandemic data, do not directly model antibody–spike protein interactions and thus fall short in predicting how existing antibody titers might target emerging or potential variants [5, 6].

To address these limitations, we utilized MINT in conjunction with the CoV-AbDab database [64] to encode antibodies—both naturally produced and vaccine-induced—and the SARS-CoV-2 spike proteins. The resulting embeddings were used to train an MLP model to predict whether an antibody–spike protein pair results in neutralization. Training was conducted using data from early pandemic variants (wild-type, Alpha, Beta, and others). For evaluation, we predicted cross-neutralization capabilities of antibodies targeting Omicron subvariants using a normalized score that reflects the likelihood of neutralization (Fig. 6a, (Methods Sec 5)). This approach mirrors a realistic pandemic scenario wherein existing antibody titers or designed antibodies need to be evaluated for their cross-neutralization capabilities against newly emerging variants.

**Figure 6:**
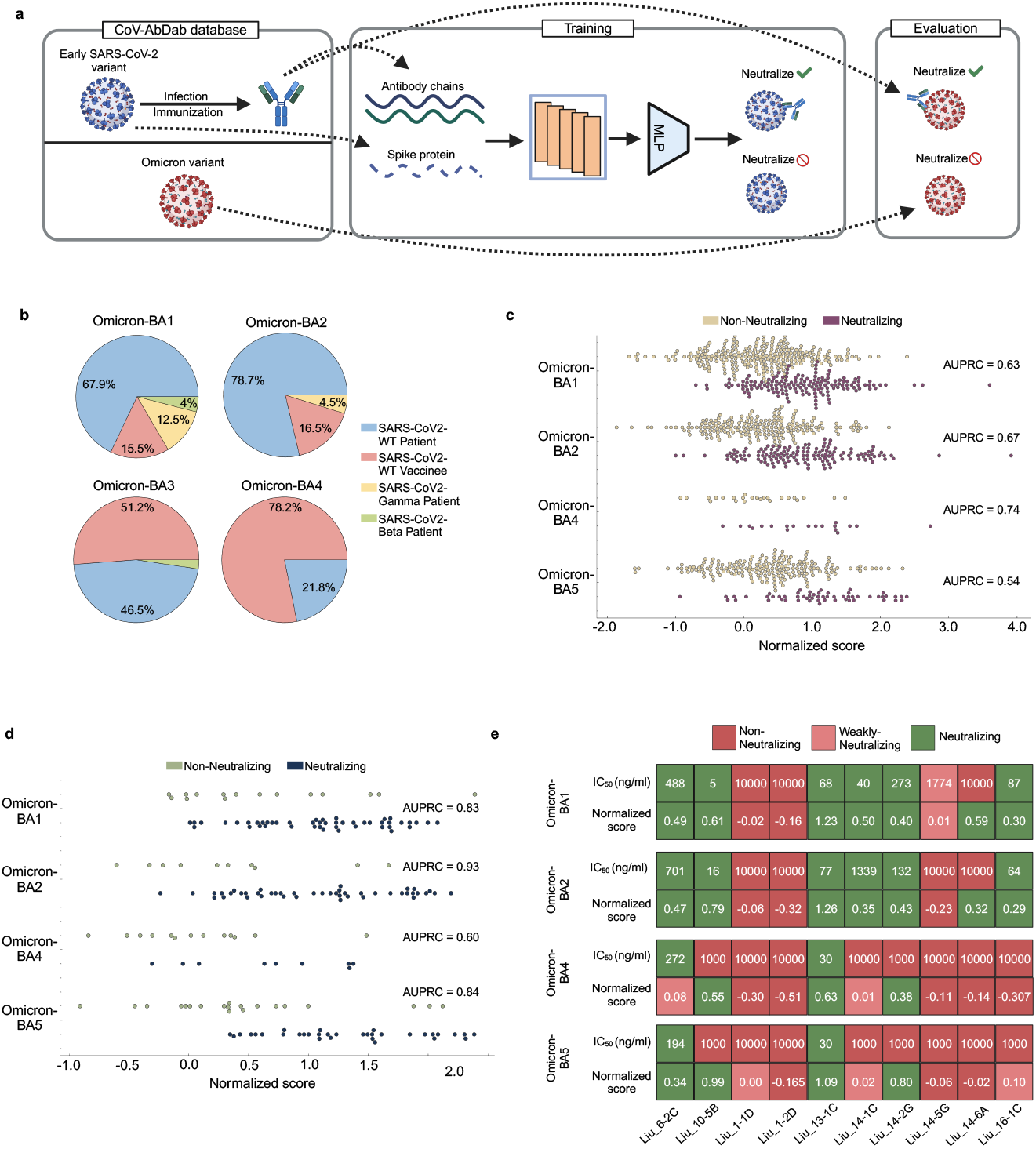
Overview and results of antibody cross-neutralization against SARS-CoV-2 variants. **a**, Procedure for predicting antibody cross-neutralization ability against SARS-CoV-2 variants. First, we extract data from the CoV-AbDab database [64] and filter entries to include antibodies produced in response to early SARS-CoV-2 variants (wild-type, Alpha, Beta, Gamma, etc) and those that target its receptor-binding domain (RBD). For evaluation, we get the entries for these antibodies against different Omicron sub-variants (BA.1, BA.2, BA.4, BA.5). The inputs to MINT are the heavy and light chain sequences along with the RBD sequence. We train an MLP on embeddings generated by MINT to predict the presence or absence of neutralization for each antibody-RBD pair. We then evaluate MINT’s performance against the Omicron sub-variants to validate the neutralization ability. **b**, The dataset composition for the constructed evaluation set across all four sub-variants of Omicron, showing the proportion of entries across different antibody origins. **c**, Distribution of the predicted normalized score for each sub-variant of Omicron across antibodies of all origin types. We group it by the actual neutralization profiles (neutralizing or non-neutralizing). We also show AUPRC values for each sub-variant. **d**, Distribution of the normalized score for each subvariant of Omicron across vaccine-induced antibodies only, grouped by actual neutralization values. **e**, Normalized score values for 10 antibodies against different variants of Omicron, along with their experimentally-derived *IC*_50_ values from Liu et al. [65]. We divide the *IC*_50_ values into non-neutralizing (=10000 ng/ml), weakly-neutralizing (*>*=1000 ng/ml and *<*10000 ng/ml) and neutralizing categories (*<*1000 ng/ml). We do the same for the predicted normalized scores: non-neutralizing (negative scores), weakly-neutralizing (positive scores less then 0.10) and neutralizing (positive scores greater than 0.10).

Fig. 6b highlights the evaluation dataset composition, showing the distribution of data points for each Omicron sub-variant and their origins, categorized as vaccine-induced or infection-derived. MINT demonstrated robust predictive performance, achieving high area under the precision-recall curve (AUPRC) values for the first three Omicron sub-variants (Fig. 6c). Notably, predictions for vaccine-induced antibodies outperformed those for infection-derived antibodies, with higher AUPRC scores observed across all variants (Fig. 6d). This enhanced performance likely reflects the greater homogeneity of vaccine-induced antibody repertoires compared to those generated through natural infection [66]. To further validate the model, we compared MINT’s normalized scores against IC50 binding affinity values reported in experimental studies. Using BBIBP-CorV vaccine-induced antibodies evaluated in Liu et al. [65], we observed an 80% hit rate for correctly identifying antibodies with consistent neutralization across Omicron sub-variants (Fig. 6e). Importantly, the model captured transitions in neutralization capacity along the evolutionary trajectory of Omicron, from BA.1 to BA.5, for antibodies such as Liu 14-5G, Liu 16-1C, and Liu 14-1C. Hence, MINT is able to discern nuanced shifts in neutralization profiles associated with variant evolution.

## Discussion

In this study, we introduced MINT, a novel protein language model designed to encode protein–protein interactions (PPIs) by simultaneously processing multiple interacting protein sequences. Unlike traditional protein language models (PLMs) that treat sequences independently, MINT incorporates cross-chain attention mechanisms to capture inter-sequence relationships. This architectural innovation preserves inter-residue context and overcomes limitations associated with previous PLMs that either concatenate embeddings or treat interacting proteins as a single continuous sequence. Our comprehensive benchmarking demonstrated that MINT significantly outperforms existing PLMs across a diverse set of supervised downstream tasks involving PPIs. In the context of antibody modeling, we showed that MINT’s training on a broad corpus of PPI data enabled it to generalize more effectively to antibody-related tasks. On TCR-epitope tasks, we highlighted MINT’s flexibility and its potential to be adapted to tasks involving protein types not present in its original training set.

We applied MINT to biologically relevant case studies to showcase its practical utility. MINT effectively modeled interacting protein sequences implicated in cancer and successfully differentiated between mutations that disrupt PPIs and those that do not. In the realm of infectious diseases, MINT accurately assessed the cross-neutralization potential of antibodies against emerging SARS-CoV-2 variants. The model’s predictions closely matched experimental data, capturing nuanced shifts in neutralization profiles associated with variant evolution and highlighting its potential in guiding vaccine design and therapeutic interventions.

Despite these advancements, there are limitations to our approach. Firstly, MINT is currently only trained on pairs of sequences from STRING using the modified MLM approach. However, our architecture can support pre-training or further finetuning on a dataset of complexes with a variable number of interacting sequences. Next, the model’s performance is inherently linked to the quality and diversity of the training data. Without finetuning, biases or gaps in PPI datasets could affect its generalizability, especially for underrepresented protein classes like peptides or interactions involving non-model organisms. For such cases, we recommend parameter-efficient fine-tuning approaches [16] that leverage domain-specific data while minimizing computational costs. These strategies can make MINT more accessible for resource-constrained research groups and enable broader impact across biomedical applications. Finally, the model’s performance on many PPI tasks is better than existing methods, but it can be improved by incorporating additional data modalities like protein structure, as proved by SaProt [33] and ESM-3 [67]. Multimodal approaches are crucial for protein representation learning because they enable models to capture complementary aspects of protein function, dynamics, and interactions that sequence-based models might overlook. Incorporating these modalities into MINT could enhance its ability to generalize to additional interaction landscapes.

MINT represents an advancement in protein language modeling by effectively capturing complex inter-sequence dependencies crucial for accurate PPI prediction. Its versatility across multiple tasks, including antibody modeling and TCR–epitope–MHC interactions, underscores its potential as a powerful tool in biomedical research. By facilitating a deeper understanding of protein interactions and their implications in disease mechanisms, MINT holds promise for accelerating therapeutic discoveries and advancing personalized medicine.

## Acknowledgments

This work was supported by the National Institute of General Medical Sciences of the National Institutes of Health under award number 1R35GM141861-01 and by a research gift from Quanta Computer. B.J. was partially supported by the Department of Energy Computational Science Graduate Fellowship under Award Number DESC0022158. S.S. was partially supported by the NSF Graduate Research Fellowship under Grant No. 2141064.

## Data availability

We retrieved physical PPI training data for MINT from STRING-DB (https://string-db.org/cgi/download) [20]. We obtained the gold-standard PPI dataset from https://figshare.com/articles/dataset/PPI_prediction from sequence gold standard dataset/21591618/3 [22], the HumanPPI dataset from https://github.com/westlake-repl/SaProt [33] and the YeastPPI dataset from PEER (https://miladeepgraphlearningproteindata.s3.us-east-2.amazonaws.com/ppidata/yeast_ppi.zip) [12]. The SKEMPI entries were downloaded from https://life.bsc.es/pid/skempi2 [23] and the PDB-Bind dataset from https://www.pdbbind-plus.org.cn/ [28]. The MutationalPPI data were obtained from https://github.com/jishnu-lab/SWING/tree/main/Data/MutInt Model [29]. The FLAB antibody datasets are available at https://github.com/Graylab/FLAb/tree/main/data [24], and the SARS-CoV-2 binding datasets at this link: https://www.biorxiv.org/content/10.1101/2020.04.03.024885v1.supplementary-material [39]. The TCR-epitope task from TDC-2 was downloaded from (https://tdcommons.ai/) [45]. The TCR-epitope-HLA data were retrieved from https://github.com/Armilius/PISTE/tree/main/data [17] and the TCR-epitope interface prediction data from https://github.com/pengxingang/TEIM [46]. We obtained experimentally validated oncoPPI data from https://github.com/ChengF-Lab/oncoPPIs [57]. Finally, we obtained SARS-CoV-2 neutralization data from https://opig.stats.ox.ac.uk/webapps/covabdab/ [64].

## Code availability

Python implementation of the MINT along with code to reproduce benchmark results and other analyses is available at https://github.com/VarunUllanat/mint.

## Authors contributions

B.J., S.S. and B.B. conceptualized the project. V.U. and B.J. constructed the training pipeline for MINT. V.U. and B.J. ran the training. V.U. performed all downstream analysis, including model benchmarking and case studies. B.B. designed and led the study. All authors contributed to writing the manuscript. We would also like to acknowledge Aditya Parekh and Anish Mudide for helpful discussions and comments.

## Online Methods

Here we describe the following methodological details: (1) curation and processing of training datasets, (2) architectural design and implementation of MINT, (3) datasets and workflows for downstream benchmarking tasks, and (4) datasets and workflows used in case studies.

### 1 STRING dataset construction

We start with 2.4 billion protein-protein interactions (PPIs) comprising 59.3 million unique protein sequences classified as physical links from the STRING database. We use mmseqs [68] to cluster the protein sequences at a 50% sequence similarity threshold, resulting in 15.6 million unique clusters. We then apply a filtering scheme that keeps only one PPI between any two clusters. This is done to ensure that PPIs between proteins belonging to the same two clusters are not repeated, so that our model can learn interactions between diverse protein types. This results in 382 million PPIs comprising 29 million unique sequences. Next, we set aside 250,000 PPIs for validation. For the training set, we apply further filtering such that no cluster in the training set is seen in the validation set. This is done to limit data leakage between the training and validation sets. Finally, we end up with 95.8 million training PPIs comprising 16.4 million unique protein sequences.

### 2 MINT architecture

Each layer of MINT receives as input an embedding of the sequences **x** *∈* ℝ^*L×F*^, a selfattention mask **m**^*att*^ *∈* ℝ^*L×L*^ and a padding mask **m**^*pad*^ *∈* ℝ^*L×L*^, where *L* is the total length of the input comprising multiple sequences and *F* is the feature dimension (*F* = 1280 in MINT). The main self-attention block we use is the MultiHeadSelfAttention used in ESM-2 [10]. We adapt this layer to return the attention matrix and the value tensor of the attention mechanism before the softmax function is applied, since we need to combine the self-attention and crossattention weights first. We present further details for the transformer block as pseudocode in Algorithm 1.

### 3 MINT training

We train MINT using the MLM objective as defined in the main text. Formally, for a given set of input amino acids in a sequence tokenized to be *x* = *x*_1_*x*_2_ … *x*_*n*_, where *x ∈* ℝ^*L×F*^, the tokens that are masked *M ∈* ℝ^*L*^, and the output vector **h** *∈* ℝ^*L×F*^, the Masked Language Modeling (MLM) loss is described as:

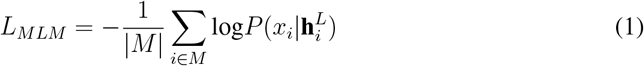

During model training, we consider 15% of tokens randomly sampled from the sequence, similar to ESM-2 [10]. For those 15% of tokens, we change the input token to a special masking token with 80% probability, a randomly chosen alternate amino acid token with 10% probability, and the original input token (i.e. no change) with 10% probability. We take the loss to be the whole batch average cross-entropy loss between the model’s predictions and the true token for these 15% of amino acid tokens.

We set the Adam optimizer with parameters *β*1 = 0.9, *β*2 = 0.98, and *ϵ* = 10^*−*8^, along with an *L*_2_ weight decay of 0.01. The learning rate was increased over the first 2,000 steps to a peak of 4*e−* 4, then gradually reduced to one-tenth of the peak value over 90% of the training. To process large proteins efficiently, we cropped long sequences to random 512-token lengths and used special BOS and EOS tokens to mark the beginning and end of proteins. We use a batch size of 2, and we accumulate gradients every 32 batches, resulting in an effective batch size of 64. The models were trained on NVIDIA A100 80GB and NVIDIA RTX A6000 GPUs in a distributed data parallel manner. We use Pytorch Lightning (https://lightning.ai/docs/pytorch/stable/) to handle the complete training and validation loops. We trained MINT for 4 million iteration steps.

#### Algorithm 1 Single functional transformer block of MINT

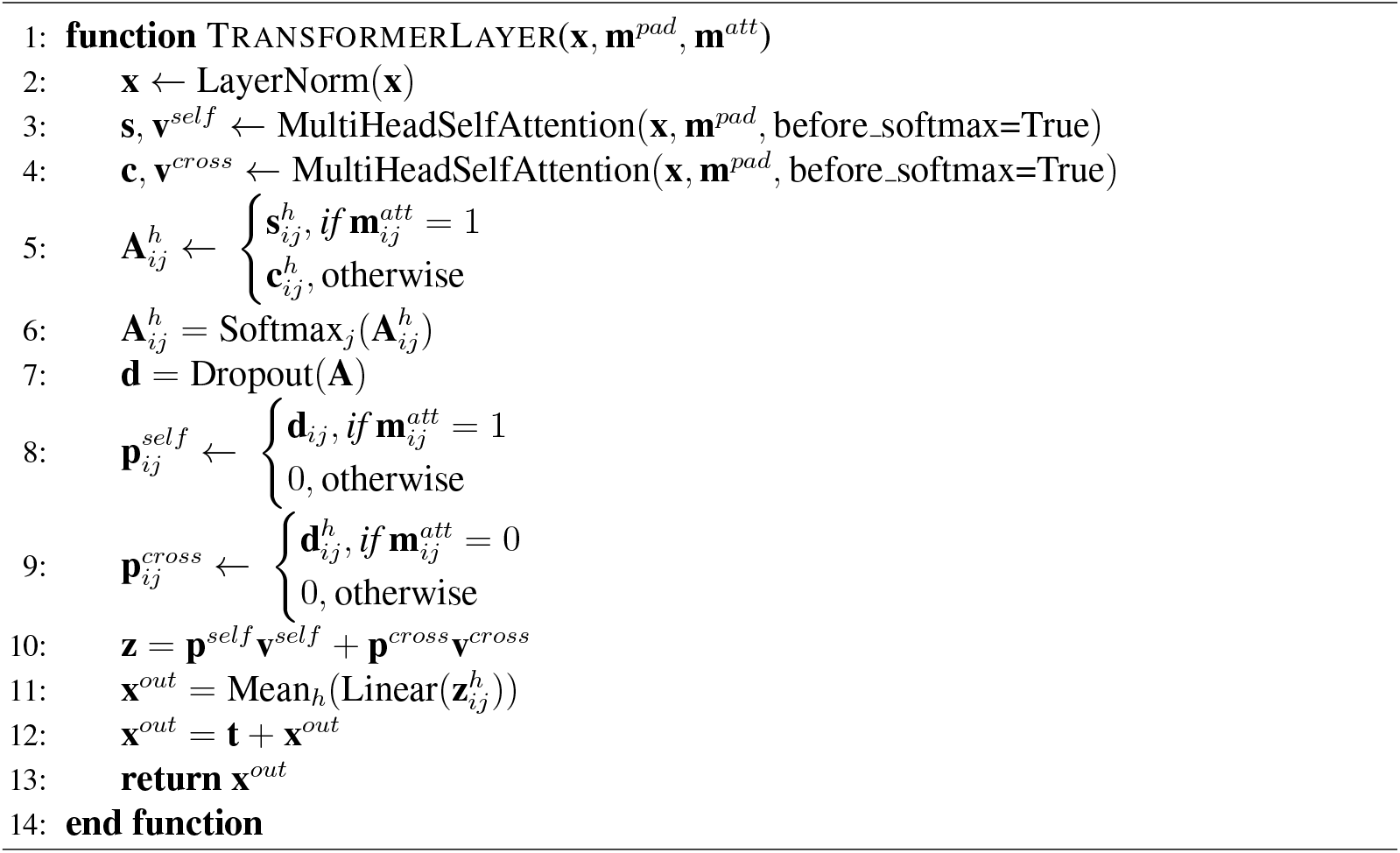

We use the perplexity metric 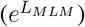 on the validation set to measure how well the model is learning across the training run. We compare this with the perplexity of ESM-2 models as a baseline on the validation set in two settings: the first where the inputs are treated as separate protein sequences and the second where the two protein sequences are concatenated. ESM-2 with single protein chains as input achieves a perplexity of 5.41 on the validation set, while concatenating achieves a perplexity of 5.16. Our MINT model achieves a validation perplexity of 4.84 at the end of its training run.

### 4 Benchmarking tasks

We compile a list of all datasets used in the across all tasks (general PPI, antibody and TCR-epitope) along with information about task type, evaluation metric used, and references in Table 1. For the antibody and TCR-epitope tasks, we use the evaluation metric used by baseline methods for consistency. For all the tasks, we use sequence-level embeddings from MINT (or a baseline PLM) to train a downstream model (MLP, ridge regression or CNN). Specifically, we extract the residue-level embedding **h** from the final layer for each group of interacting sequences, **h** *∈* ℝ^*L×F*^, where *L* is the total length of the input comprising multiple sequences and *F* is the feature dimension. We average over the sequence length *L* to get a sequence-level embedding of shape ℝ^*F*^ for each sequence. For tasks involving mutational effect prediction, we extract the sequence-level embedding for both the wild-type and mutant sequence, and use the difference between these embeddings as the input to the downstream model. The subsections describe task-specific methodologies.

**Table 1:** Summary of all datasets and tasks.

| Task | Reference | Train size | Validation size | Test size | Task | Projector used | Evaluation metric |
| --- | --- | --- | --- | --- | --- | --- | --- |
| **General PPI prediction** |  |  |  |  |  |  |  |
| Gold-standard PPI | [22] | 163019 | 59260 | 52048 | Binary classification | MLP | AUPRC |
| Human-PPI | [33] | 26319 | 234 | 180 | Binary classification | MLP | Accuracy |
| Yeast-PPI | [12] | 4945 | 95 | 394 | Binary classification | MLP | Accuracy |
| PDB-Bind | [28] | 4945 | 95 | 394 | Regression | MLP | Pearson correlation |
| SKEMPI | [23] | 4777 | - | 1929 | Regression | MLP | Pearson correlation |
| MutaionalPPI | [29] | 3406 | - | - | Binary classification | MLP | AUPRC |
| **Antibody tasks** |  |  |  |  |  |  |  |
| FLAB (Binding 422) | [36] | 422 | - | - | Regression | Ridge regression | R2 |
| FLAB (Binding 2048) | [37] | 2048 | - | - | Regression | Ridge regression | R2 |
| FLAB (Binding 4275) | [38] | 4275 | - | - | Regression | Ridge regression | R2 |
| FLAB (Expression 4275) | [38] | 4275 | - | - | Regression | Ridge regression | R2 |
| SARS-CoV2 binding | [39] | 86929 | - | - | Regression | Ridge regression | Spearman rank |
| **TCR-Epitope-MHC tasks** |  |  |  |  |  |  |  |
| TDC-Tchard | [45] | 522239 | - | 71666 | Binary classification | MLP | AUROC |
| TCR-Epitope-HLA | [17] | 28144 | 71036 | 2806 | Binary classification | MLP | AUROC |
| TCR-epitope interface prediction | [46] | 122 | - | - | Interface prediction | CNN | AUPRC |

### 4.1 General PPI tasks

As mentioned before, we evaluate two embedding strategies for the baseline PLMs. The first strategy involves making two calls to the PLM, generating the sequence-level embeddings separately, and concatenating these embeddings for all input sequences. The other involves concatenating the input sequences and generating a sequence-level embedding from the PLM.

For HumanPPI, YeastPPI and Gold-standard PPI we use the training, validation and test sets as defined in the source datasets (Table 1). For PDB-Bind and MutationalPPI we use a 10-fold cross-validation split. We split the SKEMPI dataset by protein complex to create three folds similar to previous work [69]. For all experiments except SKEMPI, we perform 3 repetitions of the training and evaluation, and report the mean and standard deviation values. For SKEMPI, we calculate the mean and standard deviation for each fold, and aggregate according to the following heuristic:

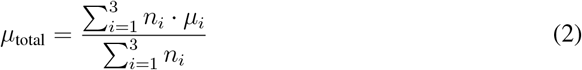

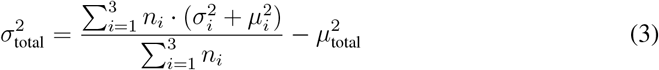

Where *n*_*i*_, *µ*_*i*_, and *σ*_*i*_ are the number of entries, mean, and standard deviation for each fold *i*.

The objective function used for training the regression tasks is the Mean Squared Error (MSE) loss, and for the binary classification tasks, we use the Binary Cross-Entropy (BCE) loss. Training was done using a 2-layer MLP with a hidden size of 640 for all experiments. We chose this number because it is half of the smallest embedding dimension across all models. We train all models for 100 epochs and choose the epoch with the best validation metric for test set evaluation. For datasets with validation splits, we use a 5-fold inner cross-validation to choose the best model.

### 4.2 Antibody tasks

#### FLAB

This dataset includes the binding energy datasets from Shanehsazzadeh et al. (2023) [36], Warszawski et al. (2019) [37] and from Koenig et al. (2017) [38], as well as the corresponding expression data from this last study. We use the sequence-level embedding over both antibody chains from MINT. We then follow [18] to perform the training experiments. This involves running a linear least squares fit with L2 regularization on the mean embedding representation features, evaluated using 10-fold cross-validation. To choose the regularization hyperparameter *λ* from the set 1, 10^*−*1^, …, 10^*−*6^, 0, we perform an additional 5-fold inner crossvalidation. We use the best performing model across each fold on the held-out testing fold and average the computed *R*^2^ across all folds.

#### SARS-CoV2 binding

We use sequences from [39] and split the data according to [19]. We use the wild-type m396 sequence from PDB (PDB ID: 2G75) and feed that into MINT separately from the mutant sequences. We extract the sequence-level embedding for both wild-type and mutant sequences, and take the difference between the two as the final embedding. We then train a ridge regression model using an *α* value of 0.01 on these embeddings.

### 4.3 TCR-Epitope-MHC tasks

We train the last layer of MINT and the downstream projector module (MLP or CNN) for the TCR-Epitope-MHC tasks.

#### TDC-Tchard

We use the ‘TDC.tchard’ dataset from TDC-2 (https://tdcommons.ai/benchmark/proteinpeptide/group/tchard/) [45], and it contains 5 different folds of train-test splits. We train the model using a batch size of 48 and an initial learning rate of 1*e*^*−*4^ for 15 epochs. We choose the best epoch as the one that has the best validation AUROC value and report those metrics. Then we take the weighted average of the AUROC across all splits (product of AUROC and test set size) to report as the final AUROC value, as suggested by TDC-2 benchmarking.

#### TCR-Epitope-HLA

For this task, we use the evaluation strategy as described in [17]. Specifically, we generate embeddings from MINT for the TCR-CDR3, HLA-CDR3, and epitope sequences, and train an MLP model for binary interaction prediction using the BCE loss. We perform three repeats and report the mean and standard deviation values in our results.

#### TCR-epitope interface prediction

We procure and use the dataset splits as constructed in [46]. For the sake of simplicity, we skip the pre-training task of TCR-epitope interaction prediction employed in the baseline model, TEIM. We also do not use the epitope autoencoder features as in TEIM. We do, however, use the same training setup after generating the embeddings for the TCR-CDR3 and epitope sequence using MINT. This entails training a three-layer CNN model to predict the contact map. We then use a BCE loss over the predicted and true contact map to train the model.

## 5 Case studies

### oncoPPI prediction

We used the dataset from [29] that contains labels on whether two human proteins continue to bind after a missense mutation as our main training set. For evaluation of mutations on oncoPPIs, we extracted the experimentally-validated mutational effect data from Cheng et al. [57] and retrieved the sequence data from UniProt. Following Cheng et al., [57], we converted the Y2H score (range of 0-4) to a 0 (non-binding) or 1 (binding) based on a cutoff of 2. For example, a Y2H score of 4 (yeast colony grows) for a mutated oncoPPI would map to a 1 (binding conserved).

The MLP is trained on the difference between wild-type and mutated PPI embeddings from MINT (in a fashion similar to 2b). Since the training dataset is very small, we performed 100 repetitions of a training and evaluation run. We then calculated a binding score *S*_*binding*_ that is defined as:

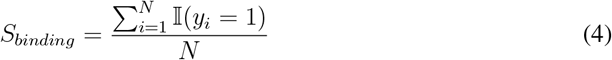

where *y*_*i*_ is the output of the model for entry *i* in the evaluation set, and *N* = 100 repetitions.

To determine an optimal threshold for classification using only the predicted binding score, we fitted the data using a Gaussian Mixture Model (GMM). This method assumes that the scores belong to two distinct groups (binding vs non-binding) and models them as a mixture of two Gaussian distributions. The threshold for classification was identified as the point where the two Gaussian distributions overlap most significantly. In our case, where the two distributions have similar variance, the threshold is approximated as the midpoint between their means, resulting in a value of 0.68. Hence, binding scores below 0.68 would be classified as non-binding, and those above as binding. We provide the AUROC curves for this analysis in Supplementary Fig. S4.

### SARS-CoV-2 Cross-neutralization prediction

We first download the entire CoV-AbDab database [64]. We then filter out non-antibody entries and entries that do not have a sequence associated with them. We then keep antibodies that fall into one of these origins: ‘B-cells SARS-CoV2 WT’, ‘Convalescent Patient (Unvaccinated)’, ‘B-cells; SARS-CoV1 Human Patient; SARS-CoV2 Vaccinee’, ‘B-cells; SARS-CoV2 WT Convalescent Patients’, ‘B-cells; SARS-CoV2 WT Vaccinee (BBIBP-CoV)’, ‘B-cells; SARS-CoV2 WT Vaccinee’, ‘B-cells; SARS-CoV2 WT Human Patient’, ‘B-cells; Unvaccinated SARS-CoV2 WT Human Patient’, ‘B-cells; SARS-CoV2 Gamma Human Patient’, ‘B-cells; SARS-CoV1 Human Patient’, ‘Bcells (SARS-CoV2 Beta Human Patient). Next, we only keep antibodies that bind to the RBD of the SARS-CoV-2 spike protein. Finally, we split the data to ensure that only neutralization values against early variants (Wild-type, Alpha, Beta, Delta, Epsilon, Gamma, Eta, Iota, Lambda, and Kappa) are seen in the training set. The evaluation set contains only Omicron sub-variants, namely BA.1, BA.2, BA.4, and BA.5. This results in 3365 data points in the training set and 1052 entries for evaluation. We download the spike protein sequences for all included variants and sub-variants from Expasy (https://viralzone.expasy.org/9556).

During training, we generate sequence-level embeddings from MINT for the antibody heavy-chain, light-chain, and the RBD domain of the spike protein. We train an MLP on these embeddings to predict whether the antibody is able to neutralize that spike protein using the BCE loss. In the training set, antibodies may be repeated since they have neutralization data against multiple variants. After training, we test the model on the evaluation set and record the raw outputs (without sigmoid activation). Since the raw outputs across the different subvariants may have different scales, we use quantile normalization to normalize the score across the sub-variants to calculate the ‘Normalized score’. We retrieve the IC50 values from [65].

